# Iterative Refinement for Structural Optimization of De Novo Protein Sequences

**DOI:** 10.64898/2026.10.27.762198

**Authors:** Chance Gammill, Lakshman Tamil

## Abstract

In this paper, we introduce an iterative refinement procedure that substitutes individual amino acid residues in a predicted protein sequence to minimize structural error; substitutions are guided by a weighted composite score of global evaluation metrics including the TM-score, GDT-TS, and RMSD. Starting from an initial sequence prediction produced by a structure-aware graph neural network (GNN) [5], the refinement procedure evaluates candidate amino acid substitutions at each position by folding the modified sequence with ESMFold [9] and accepting substitutions that satisfy both a global structural improvement criterion and a local displacement threshold. Evaluated on a matched sample of 224 proteins, this set of methods reduced mean RMSD from 8.097 Å to 4.570 Å relative to the model’s initial prediction, a 43.6% reduction, with corresponding reductions of 46.2% relative to ProteinMPNN and 27.2% relative to native ESMFolded sequences; these differences were statistically significant by paired *t*-test (*p <* 0.001). Our results show that post-prediction iterative refinement can significantly improve the structural accuracy of de novo protein sequences.

## 1. Introduction

AlphaFold [7] was a major breakthrough in the use of artificial intelligence for predicting protein structures. The resultant model for protein structure prediction in the 2020 Critical Assessment of Structure Prediction (CASP14) [12] validated an ongoing theory. Artificial intelligence models can predict the folding behavior of amino acid sequences. As protein-folding models have advanced, it is now possible to quickly evaluate the quality and structure of protein sequences, barring size limitations. This progress spurred research towards inverse protein folding, enabling researchers to generate a protein sequence that folds into their desired protein structure. However, each model has its own limitations, so we designed a method that uses both a protein folding model [9] and an inverse protein folding model [5], allowing them to benefit from each other through iterative refinement for post-prediction optimization. Although an inverse protein folding model can quickly predict a structure close to the desired conformation, slight deviations can cause major problems for drug development and lead to adverse side effects. By spending more time optimizing the initial prediction using feedback from the protein folding model, we can significantly reduce the inverse protein folding model’s structural folding error, making generated de novo protein sequences more structurally accurate at the cost of time and compute. This approach of using logit information from the inverse protein folding model to optimally test the different protein sequences and only continuing with the best-performing sequence is what we have titled: Iterative Refinement.

## 2. Data Accessibility

The evaluation data used in this paper is identical to the data described in our companion paper [5] and is summarized below. Protein crystallographies were obtained from the Protein Data Bank [2] in mmCIF format with the AlphaFold3 [1] `fetch_databases.sh` utility command, in February 2026. The structures were decomposed into individual chains and converted to backbone coordinate representations of the atomic positions N, C*α*, C, and O. All chains were utilized regardless of native length and missing residues, then truncated to a fixed 128 residues; short chains were padded with a placeholder token “unknown residue”. No filtering by resolution, experimental method, or chain type was used, and the gaps in the deposited coordinates were passed through unchanged.

Because truncation, padding, and chain selection were performed without regard to whether the resulting fragment could fold in isolation, some evaluation targets may be structurally unrealizable. These implausible configurations were retained, and each method was required to approximate them as closely as possible. This directly affects the performance of sequence recovery methods and explains ProteinMPNN [4] underperforming relative to its published sequence-recovery capabilities.

This work does not train any model and only uses existing models; we use a GNN model [5] as a generator and ESMFold [9] as an external structure predictor. Any overlap between ESM-Fold’s pretraining data and the evaluation data cannot be accounted for and is acknowledged as a limitation of relying on a pretrained predictor for evaluation. Of the 10,000 evaluation structures, a matched comparison was performed for 224 targets across all four methods (native, initial prediction, refined, and ProteinMPNN); a practical limit set by the cost of running the complete refinement and ProteinMPNN pipelines per target.

## 3. Methods

Our approach consists of three stages: an initial sequence is generated for a target backbone by a structure-aware graph neural network (GNN) [13] (Section 3.1); the resulting structure, together with the baseline structures produced by ProteinMPNN, is evaluated using a set of structural comparison metrics (Section 3.2); and the initial GNN sequence is then optimized through an iterative refinement procedure that searches for per-position substitutions improving structural agreement with the target backbone (Section 4).

### 3.1. Initial Sequence Model

The structure-aware graph neural network described in our companion work [5] generates the initial predicted sequence for each target backbone, as summarized here.

The model adapts the ProteinMPNN [4] encoder–decoder architecture, using four bidirectional message-passing [6] encoder layers and four decoder layers with node, edge, and hidden feature dimensions of 128, over a structural *k*-nearest-neighbor graph [3] with *k* = 20. Unlike the ProteinMPNN decoder, the model described in our companion paper is fully sequential and soft auto-regressive, meaning that each position is decoded using probability-weighted embeddings of previously decoded positions. This does two important things. Firstly, the output at each position is a categorical probability distribution **p**_*i*_ over the 20 standard amino acids, which iterative refinement can use to rank candidate substitutions (Section 4). Secondly, the model was trained for structural optimization rather than native sequence recovery, so **p**_*i*_ (the logits) are biased toward structurally preferred amino acids, rather than sequentially typical amino acids.

### 3.2. Evaluation Metrics

We evaluated the structural accuracy of the predicted and native sequences by folding the sequences using ESMFold [9] to obtain a predicted three-dimensional coordinate structure, which was then compared against the ground-truth backbone structures. All structural comparisons were performed after optimal superposition using the Kabsch algorithm [8]. The following metrics were used to create scores for each evaluation, and the weight of each metric is decided by the user before refinement, allowing personalized optimization of their protein sequence, creating a final score per prediction that is used to compare each created sequence and allows for adjustable comparisons of de novo proteins, without a “true” sequence.

Structure prediction used ESMFold v1 (`facebook/esmfold_v1`) via the Hugging Face `transformers` library (`EsmForProteinFolding`, `EsmTokenizer`), run in full float32 precision rather than half precision for numerical stability, frozen throughout, with 4 recycles. As a single sequence structure predictor, ESMFold does not use multiple sequence alignments or structural templates. An explicit random seed was not set for ESMFold inference itself, as structure prediction is deterministic given the input sequence; stochasticity in the pipeline was introduced through sampling-based sequence generation (Section 4). Positions padded with the placeholder “unknown residue” token were excluded from the sequence passed to ESMFold rather than folded as an explicit unknown residue token; ESMFold therefore only ever folds the unpadded residues of each target. However, these excluded positions are still represented in the per-residue error matrices and heatmaps (Figures 9) as placeholder columns aligned to the fixed 128-residue representation, as the dimensions of the matrices are kept uniform across all evaluated proteins regardless of native length.

Both the initial GNN and ProteinMPNN predictions were evaluated in a matched best-of-5 sampling budget: five sequences per target were generated for each method, and the sequence with the lowest self-reported RMSD for each model was carried forward for structural comparison. ProteinMPNN sequences were generated using the default model weights (`v_48_020`, vanilla non-soluble non-CA only weights, 48 edges, 0.20 Å training noise), with `num_seq_per_target=5`, sampling temperature 0.1, zero backbone noise, and a fixed random seed of 42, applied identically throughout the evaluation target set.

Iterative refinement was applied as a separate single run starting from one initial GNN prediction per target, rather than to the best of the five candidates; iterative refinement’s two-pass candidate search (Section 4) consumes a significant amount of per-target computational effort, causing refinement and the best of 5 baselines to not be matched in computational expense.

TM-score, RMSD, and GDT-TS were computed via a custom inline implementation following the reference TM-score definition of Zhang and Skolnick [16], using a singular value decomposition-based Kabsch alignment. The length dependent normalization term was computed as *d*_0_ = max(0.5, 1.24 (*L*_target_ − 15)^1*/*3^ − 1.8) for *L*_target_ *>* 15, and *d*_0_ = 0.5 otherwise, providing an explicit short chain correction. GDT-TS used the same Kabsch alignment as TM-score and RMSD.

The metrics defined in Section 3.2.1 quantify structural agreement when reporting results, and are combined into the composite score that dictates candidate acceptance during refinement (Equation 5). Their definitions are therefore provided in full here rather than by reference. The use and definitions of these metrics follow the definitions used in our companion work [5].

#### 3.2.1 Global Structural Metrics

##### Root-Mean-Square Deviation (RMSD)

We computed RMSD [11] between C*α*s using the ESMFold-predicted structure of the predicted sequence and the ground-truth backbone after Kabsch superposition, as given in Equation 1:

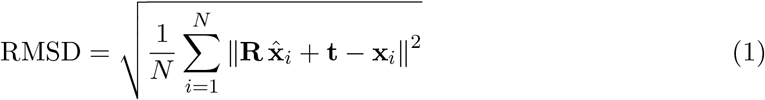

where 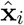 and **x**_*i*_ denote the predicted and native C*α* coordinates of residue *i*, respectively, **R** and **t** are the optimal rotation and translation obtained via Kabsch alignment, and *N* is the number of residues included in the comparison.

##### TM-score

The template modeling score (TM-score) [16] is a length-normalized measure of structural similarity that is less sensitive to local outliers than RMSD, as given in Equation 2:

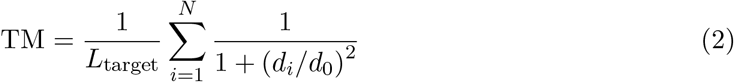

where *d*_*i*_ is the distance between the *i*-th pair of aligned C*α* atoms after superposition, *N* is the number of aligned residues, and 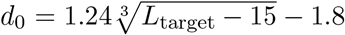 is a length dependent normalization factor. Statistical analysis of PDB structures indicates that scores below approximately 0.17 are characteristic of randomly chosen unrelated protein pairs, whereas scores above 0.5 generally correspond to pairs sharing the same fold in SCOP/CATH [14].

##### GDT-TS

The Global Distance Test–Total Score (GDT-TS) [15] measures the fraction of residues whose C*α* atoms fall within multiple distance thresholds after Kabsch alignment, as given in Equation 3:

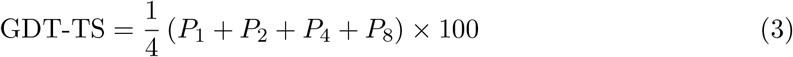

where *P*_*d*_ denotes the percentage of residues whose C*α* atoms are within *d* Å of their native positions.

##### pLDDT

The predicted Local Distance Difference Test (pLDDT) [10] reported by ESMFold provides a per-residue estimate of model confidence in the predicted local backbone geometry. Higher pLDDT values indicate greater confidence in the accuracy of local structural features, while lower values typically correspond to flexible or poorly resolved regions. Although high scores correlate with the ESMFold model’s folding confidence for that sequence, in de novo proteins a lower pLDDT is not necessarily a negative result, as it only reflects the model’s confidence in that prediction. This reflects a lack of knowledge about that amino acid sequence, which is an expected consequence of our approach; the pLDDT values obtained for each sequence type are reported in Section 5.2.

## 4. Iterative Refinement

To improve the structural quality of a model’s predicted sequence, starting from an initial sequence prediction, the procedure performs two passes over the sequence, summarized in Section 4.1. It tests candidate amino acid substitutions at each position and accepts changes that improve structural accuracy. This procedure is designed to avoid the disruption of inter-residue dependencies. Substitutions accepted in earlier predicted positions will influence the predictions of later residues; the second pass allows positions that were initially skipped or sub-optimally predicted to benefit from the updated sequence context.

### 4.1. Procedure Overview

The complete procedure is summarized in Figure 1. Starting from the initial sequence *S*^(0)^ and the probability distributions per position**p**_*i*_ produced by the design model (Section 3.1), each pass visits positions in randomized order. At position *i*, the top-*k* residues under **p**_*i*_ form the candidate set *C*_*i*_, with *k* = 5 in the first pass and *k* = 20 in the second; each candidate substitution is folded with ESMFold and scored. A substitution is accepted only when it both improves the weighted global objective (Equation 5) and produces a local displacement below *e*_thresh_ = 0.1 Å (Equation 4); among the surviving candidates, the one which minimizes 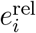 is selected (Equation 6). Positions at which no candidate satisfies both conditions retain their original residue and are re-evaluated in the following pass. Candidate selection is described in Section 4.2 and the acceptance test in Section 4.3.

**Figure 1:**
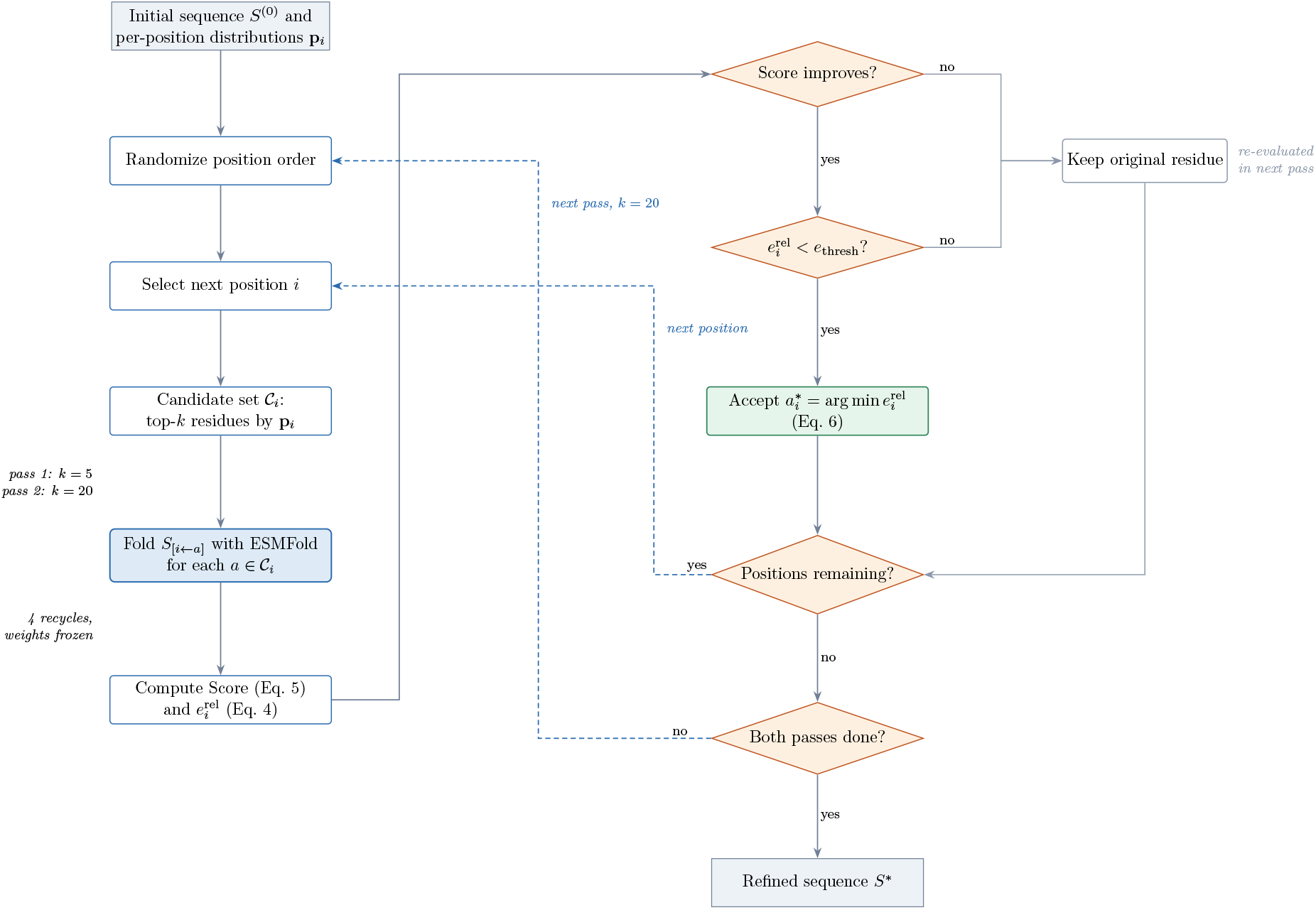
The two-pass iterative refinement procedure, from the initial sequence *S*^(0)^ to the refined sequence *S*^∗^, with dashed blue paths marking the loop over positions within a pass and the loop over the two passes.

### 4.2. Candidate Selection and Ranking

At each position *i*, the model creates a probability distribution **p**_*i*_ over the 20 standard amino acids. The candidates are ranked according to their predicted probability, and iterative refinement evaluation is limited to the top-*k* candidates per pass. During the first pass, only the top *k* = 5 candidates are considered. This limitation restricts substitutions and limits model divergence, preserving structural motifs and bonding preferences encoded in the model’s learned probability distribution. This reduces the risk of disrupting well-predicted regions and selects the best “reasonable” choice. In the second pass, the full set of 20 amino acids is considered at each position, allowing more exploration once the sequence has already been partially stabilized in the first pass.

The decoding order during each pass is randomized, meaning that no position is treated as a fixed anchor. This avoids the positional bias that would arise from always beginning substitutions at the terminus, and allows the refinement to converge to solutions that are not constrained by the order in which residues happen to be evaluated.

### 4.3. Acceptance Criterion and Convergence

The local relative displacement error at position *i* for a candidate substitution *a* is defined as the distance the ESMFold-predicted C*α* at position *i* moves when the substitution is applied:

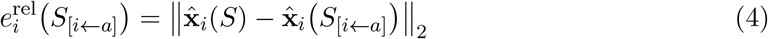

where 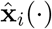 denotes the ESMFold-predicted C*α* coordinate at position *i* for the given sequence.

A substitution at position *i* is accepted only if two conditions are met. (1) Firstly, the substitution must produce a global improvement, dictated by a weighted composite of GDT-TS, TM-score, and RMSD (Equation 5). (2) Secondly, the local relative displacement error at position *i* falls below the acceptance threshold *e*_thresh_ = 0.1 Å. Condition (2) is enforced regardless of the improvement in condition (1); a substitution that improves the global score but produces a local displacement at or above *e*_thresh_ is discarded.

The composite global objective used to evaluate condition (1) is a weighted sum of the TM-score, GDT-TS, and the RMSD score:

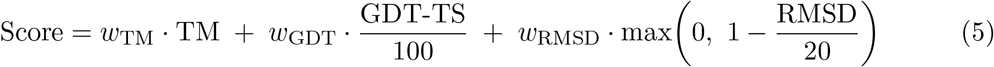

where TM and GDT-TS are calculated as in Equations 2 and 3, and RMSD is bounded to the [0, 1] range by clipping at 20 Å before weighting. All three terms are constructed so that higher values indicate better structural agreement, and the weighted sum is maximized. The weights used to generate the results reported in this manuscript were *w*_TM_ = 0.4, *w*_GDT_ = 0.3, and *w*_RMSD_ = 0.3.

The dual requirement system of global improvement and local error below *e*_thresh_, drawn as the two decision stages of Figure 1, prevents iterative refinement from greedily accepting substitutions that reduce global error at the cost of disrupting the local fold. Although theoretically this procedure could enforce hard convergence over an infinite number of passes, the time for convergence can be long, and each pass achieves diminishing returns. So, for the sake of testing, it has been limited to 2 passes.

Formally, among the candidates in *C*_*i*_ that satisfy both acceptance conditions, the selected substitution is the one with the minimum local relative displacement error:

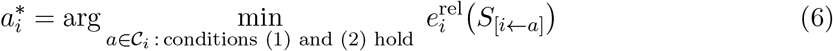

where *C*_*i*_ is the set of top-*k* candidates at position *i* for the current pass, and *S*_[*i*←*a*]_ denotes the sequence obtained by substituting position *i* with candidate *a* while all other positions remain unchanged. If no candidate at position *i* satisfies both acceptance conditions during a pass, the original residue at that position is left unchanged and is re-evaluated as a candidate site in the subsequent pass.

## 5. Results

We evaluated the iterative refinement procedure on a separate set of *n* = 224 evaluation proteins, not included in the training data, comparing ESMFold-predicted structures for four sequence categories: Native, Predicted, Refined, and ProteinMPNN, against the provided crystal structure coordinates after Kabsch-superposition. The best-of-5 sampling budget applied to the initial GNN and ProteinMPNN predictions is described in Section 3.2, and the two-pass search budget used by refinement in Section 4. Of the 224 evaluated proteins, 221 had complete and valid results across all four methods and were used for the paired statistical comparisons reported below. The remaining three proteins were excluded due to missing results, most likely through degenerate or empty sequences, causing downstream folding or scoring failure.

### 5.1. Evaluation Metrics

The three primary structural metrics reported here are defined in Equations 1, 2, and 3 of Section 3.2.1. Our model with iterative refinement achieved a mean RMSD of 4.570 Å (Figure 2), a significant improvement over ProteinMPNN and the native sequence as assessed by ESMFold: a difference of 3.930 Å when compared against ProteinMPNN (a 46.2% reduction) and 1.710 Å when compared against the native sequence a 27.2% reduction. Our model also achieved an average TM-score of 0.675, a 36.4% increase over ProteinMPNN and an 11.4% increase over the native sequence. Additionally, it obtained a GDT-TS of 61.775, a 44.8% increase over ProteinMPNN and a 14.0% increase over the native sequence. However, this improvement comes at a cost. The iterative refinement procedure optimizes solely for structural proximity to the target backbone, without penalizing deviations from the native sequence. As a result, sequence accuracy drops substantially between the native sequence and the refined output.

**Figure 2:**
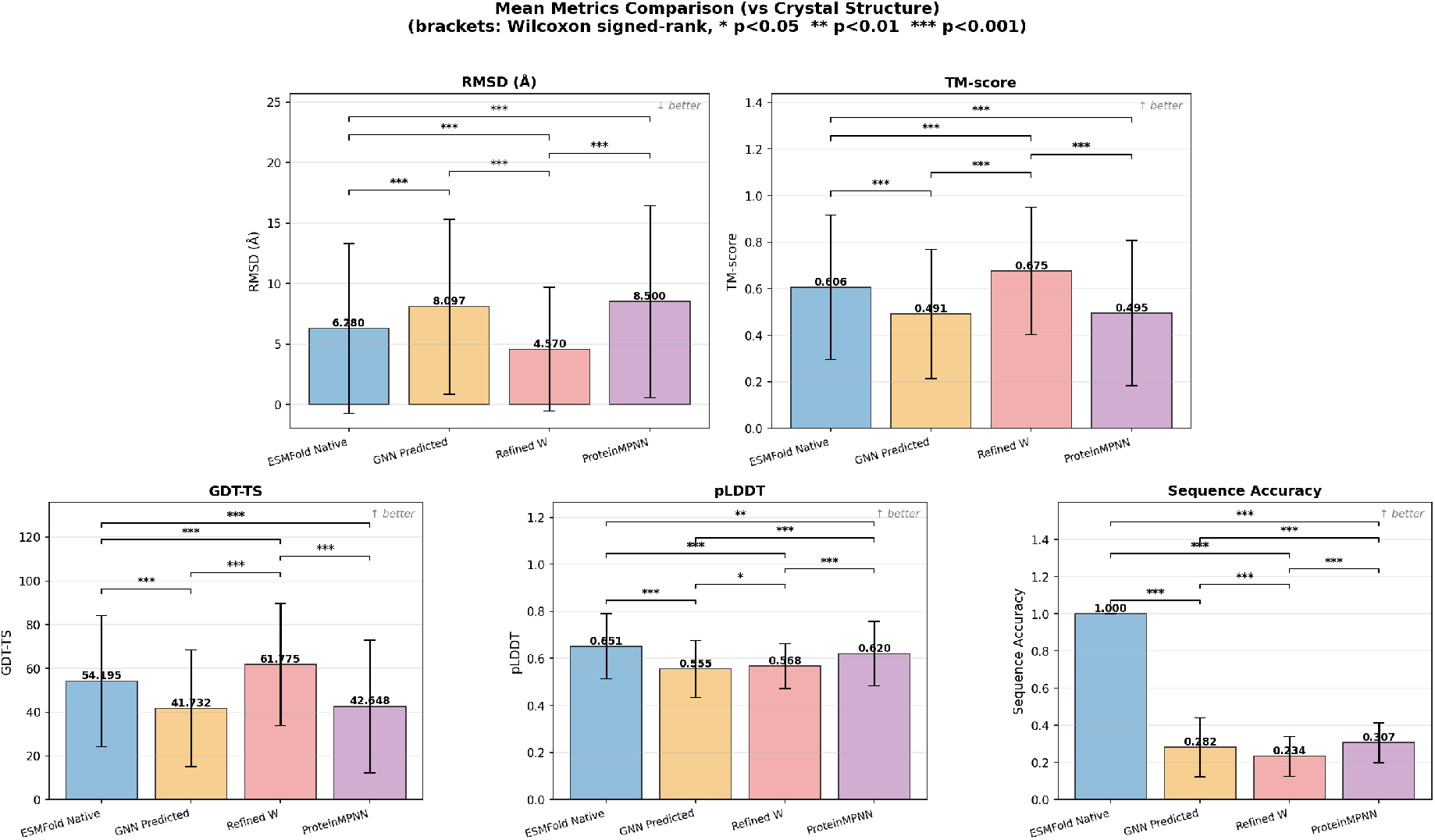
Mean structural comparison metrics across all evaluated proteins, four sequence types are compared: the PDB native sequence, our model’s initial prediction, the sequence after iterative refinement, and ProteinMPNN’s predicted sequence.

**Figure 3:**
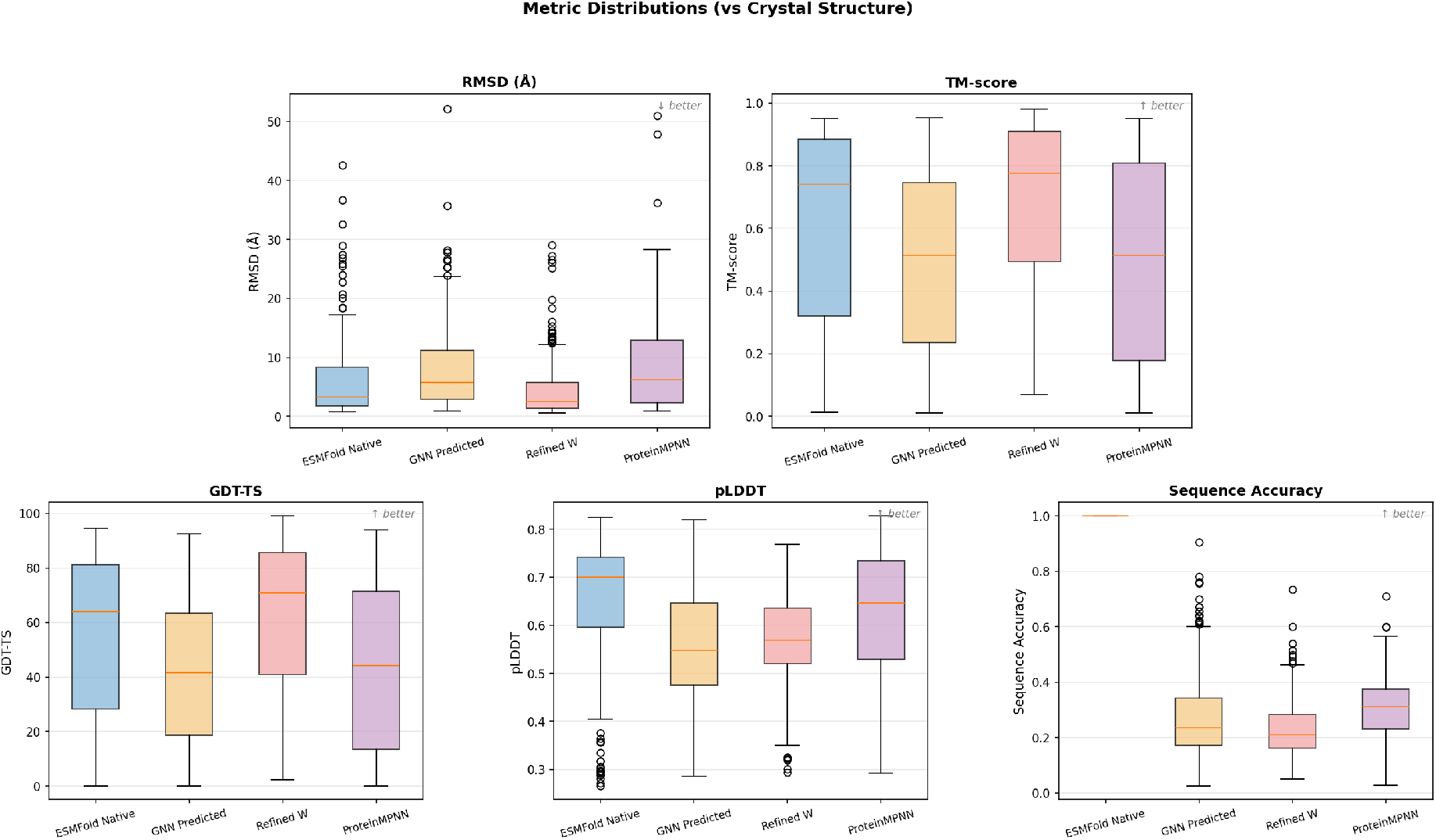
Distribution of structural comparison metrics across all evaluated proteins, shown as box plots giving the median, inter-quartile range, and spread for each metric. Four sequence types are compared: the PDB native sequence, our model’s initial prediction, the sequence after iterative refinement, and ProteinMPNN’s predicted sequence.

Because all four methods were evaluated on the identical set of *n* = 221 proteins with complete results, paired comparisons are possible. Paired *t*-tests (confirmed by Wilcoxon signed-rank tests) showed that refinement produced statistically significant improvements over every baseline across all primary structural metrics (*p <* 0.001 for RMSD, TM-score, and GDT-TS, refined vs. initial prediction, vs. native, and vs. ProteinMPNN). Notably, the initial unrefined GNN prediction and ProteinMPNN were *not* significantly different from one another on RMSD (*p* = 0.221), TM-score (*p* = 0.805), or GDT-TS (*p* = 0.516), indicating that the initial sequence model performs comparably to ProteinMPNN on this truncated evaluation set, and that the significant structural improvement reported here is attributable specifically to the iterative refinement procedure rather than to the base sequence model.

The aggregate improvement reported above spans a wide range of per-protein outcomes. Figure 4 shows 7SCA, chain AJ, a case where refinement produced a large improvement over the initial prediction; Figure 5 shows 7SER, chain A, a case representative of the median outcome across the evaluation set; and Figure 6 shows 7S94, chain E, one of the exceptions in which ProteinMPNN outperformed the refined sequence.

**Figure 4:**
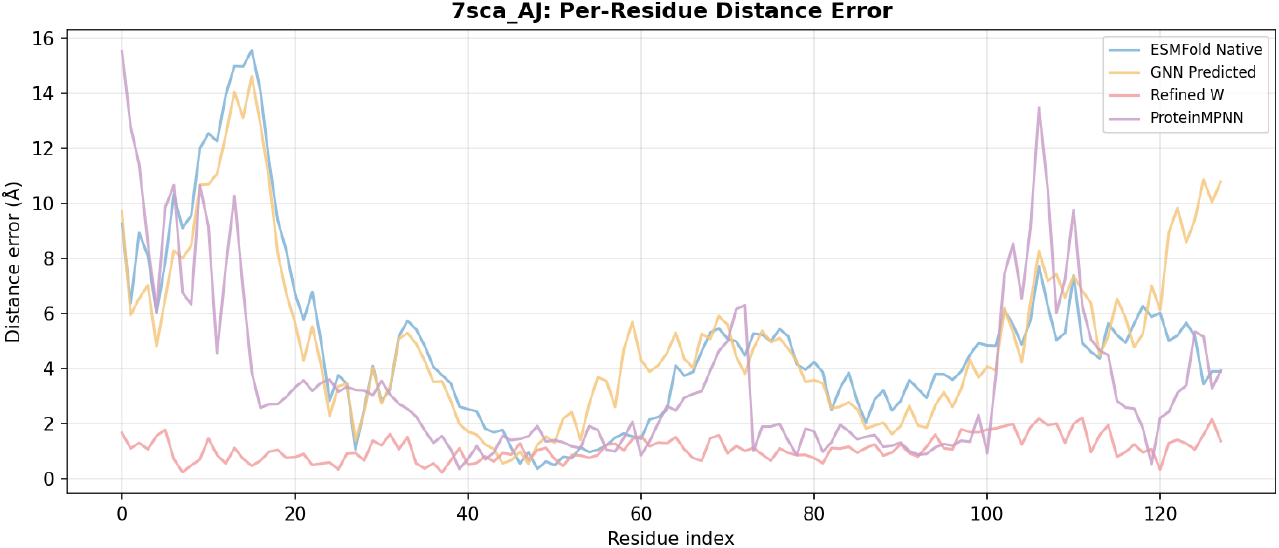
7SCA AJ is a case of large improvement from iterative refinement.

**Figure 5:**
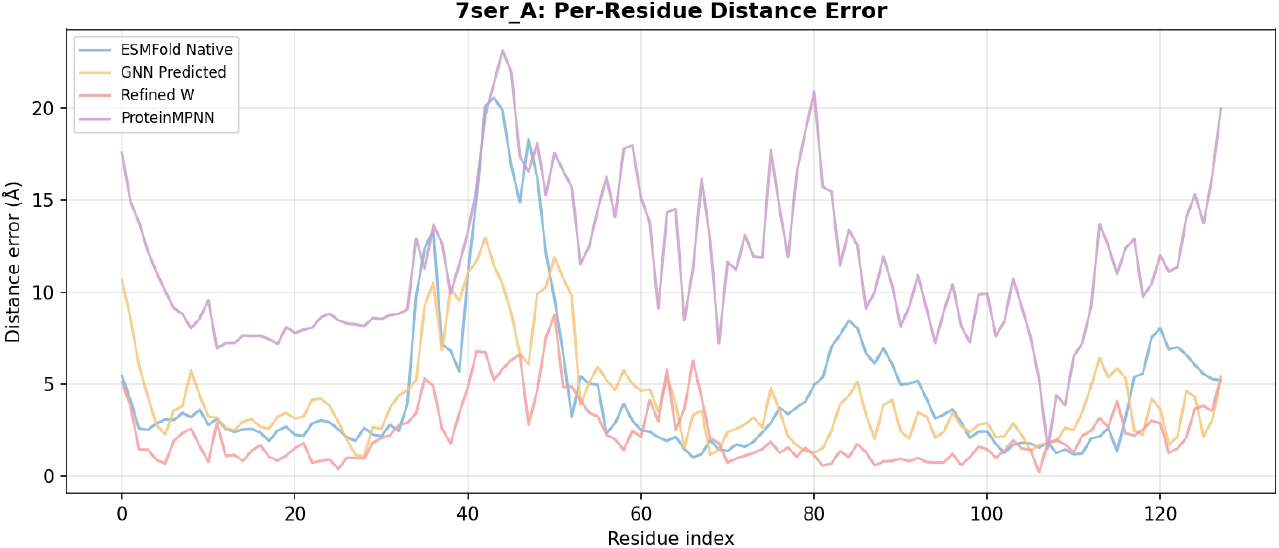
7SER A is a case representative of the median outcome across the full evaluation set.

**Figure 6:**
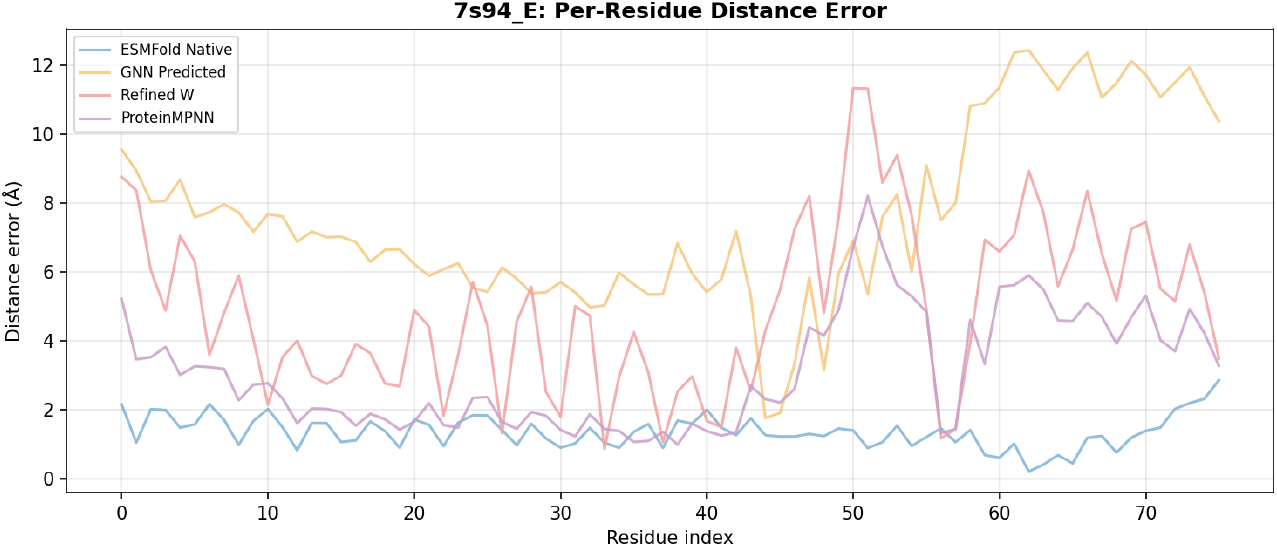
7S94 E is a case where ProteinMPNN outperformed the refined sequence.

### 5.2. pLDDT

The refined and unrefined predicted sequences achieved mean pLDDTs of 0.568 and 0.555, respectively. This difference is statistically significant (paired *t*-test *p* = 0.009), indicating that iterative refinement slightly improves ESMFold’s confidence relative to the initial prediction, although the magnitude of the change is small.

The mean pLDDTs of the refined and unrefined predicted sequences remained well below ProteinMPNN (0.620) and the native sequence (0.651). This outcome was expected, as more well-known and natural sequences yield higher pLDDTs. So, lower pLDDT scores will correlate with a divergence of protein sequences from the natural behavior of proteins. However, this also raises concerns about reduced predictability, possible adverse structural effects, and structural collapse. This unpredictability is an inherent issue with this approach, and if necessary, it can be accounted for in the composite global score.

### 5.3. Limitations and consequences of shortened chains

In Figure 3, it can be seen that even with iterative refinement, not all sequences can be optimized to a reasonable degree; this follows from the fixed 128-residue truncation described in Section 2. This is most likely caused by the protein’s tertiary structure, which falls outside of the truncated amino acid sequence, so it is possible that no shortened sequence would be capable of reproducing the provided protein structure. It is also the reason that ProteinMPNN exhibits significantly worse results than it would typically score. However, it also shows that there are limitations to some of the top-performing models, as the model after iterative refinement is able to optimize on previously unpredictable structures. The per-residue evidence for this truncation effect is presented in Section 5.4.

Figures 7 and 8 show two representative cases of this limitation directly: 7RS6, chain C, where refinement provided negligible benefit, and 7S6Q, chain E, where refinement improved structural agreement, but the refined structure remained poor in absolute terms.

**Figure 7:**
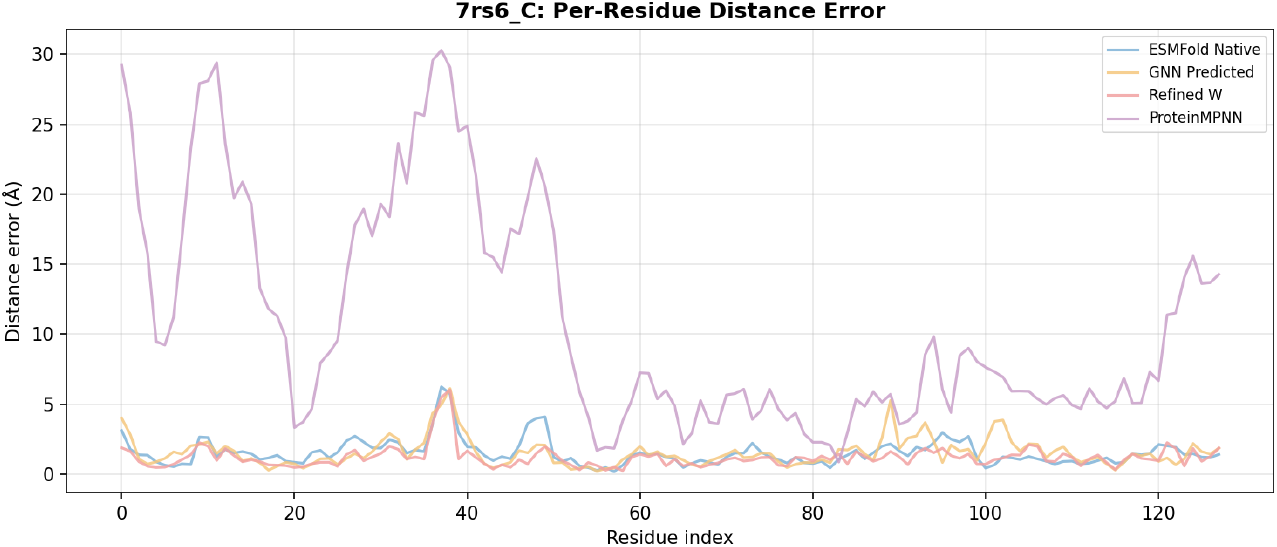
7RS6 C – refinement provided negligible benefit. Per-residue structural error (Å) by sequence position, comparing native, initial prediction, refined, and ProteinMPNN sequences against the crystal structure.

**Figure 8:**
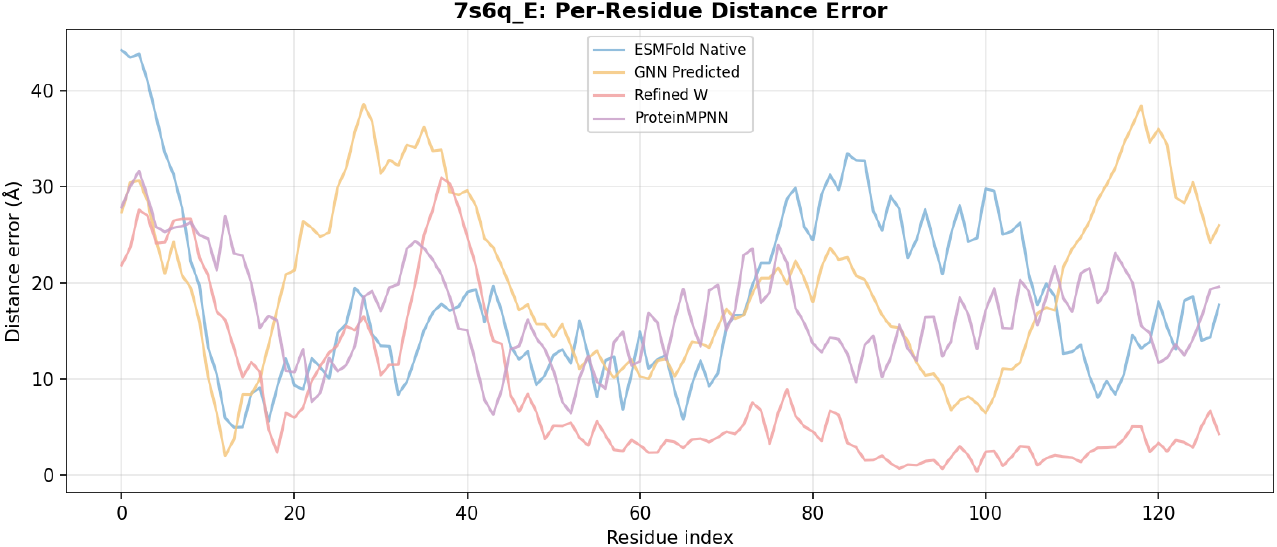
7S6Q E – refinement helped, but the refined structure remained poor in absolute terms.

### 5.4. Per-Residue Error Heatmaps

Per-residue error heatmaps aligned by sequence position (Figure 9) provide direct evidence for the truncation-driven hypothesis of Section 5.3: the error is not distributed uniformly along the sequence for any method, but instead concentrates disproportionately near the boundary of the retained 128-residue window, consistent with the loss of tertiary contacts that would otherwise stabilize those residues in the full-length native structure. This pattern is visible across all four sequence types, indicating that it is a property of the truncated set itself rather than of any single design method.

**Figure 9:**
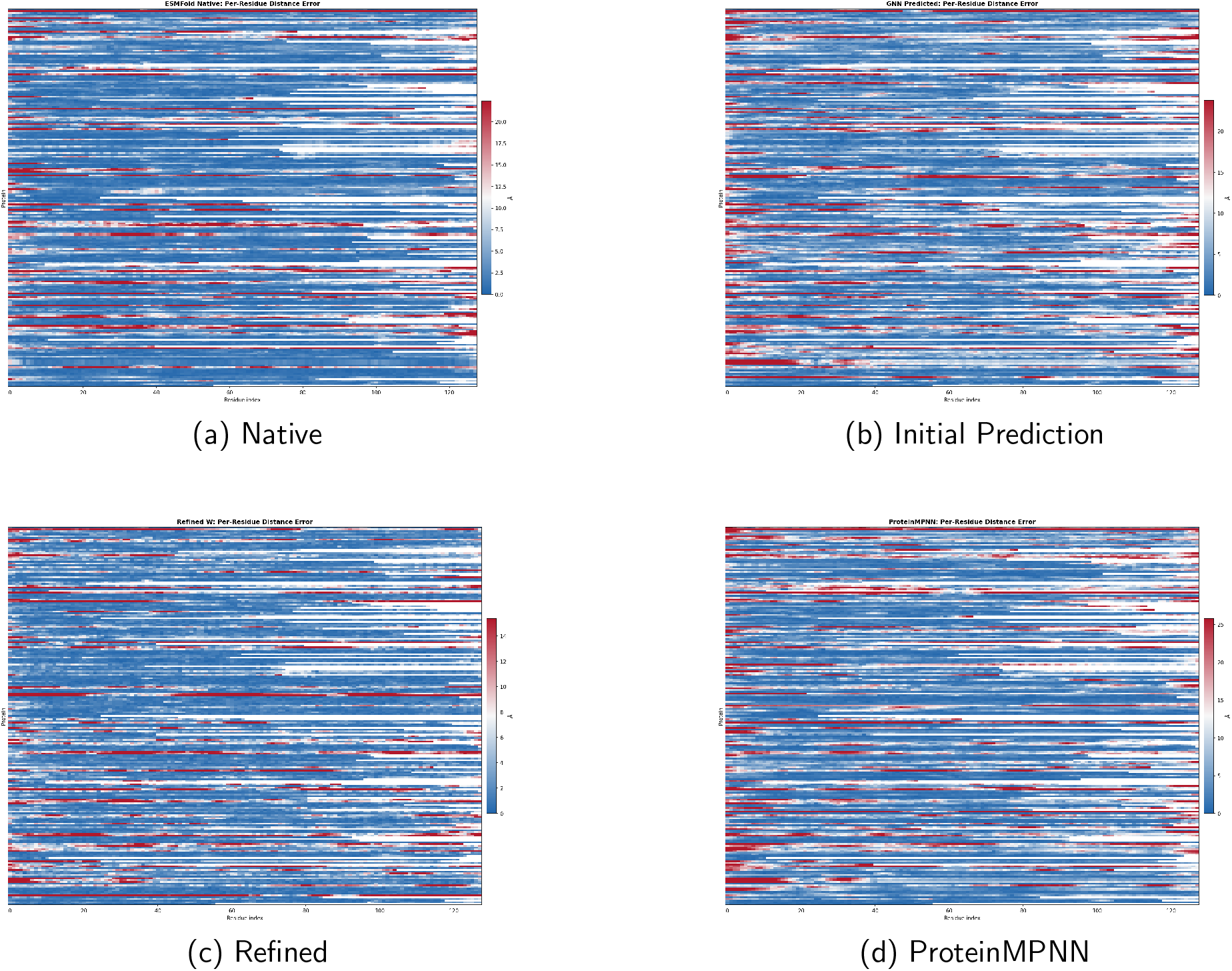
Per-residue structural error (Å) across all evaluated proteins (*n* = 224), aligned by sequence position, for each of the four sequence types.

The proportion of evaluation targets that were truncated (native length exceeding 128 residues) versus left intact was not tracked separately in this evaluation; a stratified analysis distinguishing truncated from non-truncated targets, or evaluation using intact structural domains rather than a fixed length truncation, is left to future work.

## 6. Conclusion

This study introduced an iterative refinement method (Section 4) that utilizes a model’s predicted probability distribution to identify per-position substitutions that enhance global structural metrics while maintaining local structural motifs, accepted under the dual criterion of Equations 5 and 4. This post-processing optimization reduced the mean RMSD from 8.097 Å to 4.570 Å relative to the initial model prediction, resulting in a 43.6% reduction. All structural metric improvements were statistically significant when evaluated by a paired *t*-test (*p <* 0.001) in the matched 221-protein samples (Section 5.1), demonstrating that iterative refinement can further improve protein structural accuracy. The initial unrefined GNN and ProteinMPNN predictions performed comparably, and no statistically significant differences were observed. Therefore, the significant differences reported in this paper are attributable to the refinement procedure, and not the underlying model. This finding supports the central claim that the post-prediction structural iterative refinement, rather than the initial design model, drives the observed improvement in structural accuracy. The structurally focused training of the initial model may have contributed by providing a probability distribution already biased toward structurally plausible substitutions, which may have facilitated more effective convergence during refinement than would have been possible from a sequence-oriented-recovery model. Although iterative refinement is most effective for models trained on structural accuracy, it may also perform well for protein sequence recovery models, though some structural preferences and bonding motifs may be diminished.

Some limitations should be considered. First, the iterative refinement method requires multiple ESMFold passes per amino acid, resulting in significant computational expense:

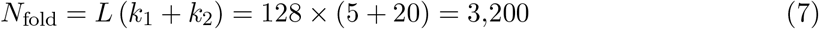

where *L* is the number of unpadded residues, and *k*_1_ = 5 and *k*_2_ = 20 are the candidate-set sizes in the first and second passes, respectively. The maximum evaluated length is *L* = 128. By Equation 7, a single protein sequence requires 3,200 forward passes, making the procedure substantially more expensive than a single-pass sequence design. Second, ESMFold is used both to guide candidate selection during refinement (Section 4.2) and to evaluate the resulting structures (Section 3.2). The reported metrics demonstrate convergence with ESMFold’s structural predictions but do not independently validate structural accuracy. Experimental testing is required to determine whether the observed improvements reflect true structural accuracy or convergence toward ESMFold-specific biases.

In conclusion, this work presents an iterative refinement procedure that directly optimizes the three-dimensional conformational accuracy of predicted protein sequences. It prioritizes structural metrics over sequence recovery and iterates over candidates dictated by the predicted probability distribution, as summarized in Figure 1. The output of the iterative refinement can be specialized for a desired structural metric. Finally, this approach achieved a 46.2% reduction in RMSD compared to a top-performing sequence recovery model. These results demonstrate that post-prediction refinement is a powerful tool for improving the structural accuracy of de novo protein design.

## 7. Declarations

### Use of generative AI

Claude (Anthropic) was used for LaTeX formatting, editing, and drafting the TikZ source for Figure 1. All figures were reviewed, corrected, and finalized by the authors. The authors assume full responsibility for the generated content. No generative AI tool was used to produce, analyze, or interpret the experimental results.

### Author contributions

C.G.: conceptualization, methodology, software, validation, formal analysis, data curation, writing — original draft. L.T.: supervision, writing – review & editing.

### Data availability

PDB accession identifiers and evaluation splits will be deposited with the archived code release.

### Code availability

Model code, evaluation scripts, trained weights, and other code are available at https://github.com/Justworkname/GenomeGenerator

### Ethics approval

Not applicable; this study used publicly available protein structures and did not involve human participants or animals.

